# Periodic and Aperiodic Neural Activity Displays Age-Dependent Changes Across Early-to-Middle Childhood

**DOI:** 10.1101/2021.10.19.464887

**Authors:** Aron T. Hill, Gillian M. Clark, Felicity J. Bigelow, Jarrad A. G. Lum, Peter G. Enticott

## Abstract

The neurodevelopmental period spanning early-to-middle childhood represents a time of significant growth and reorganisation throughout the cortex. Such changes are critical for the emergence and maturation of a range of social and cognitive processes. Here, we utilised both eyes open and eyes closed resting-state electroencephalography (EEG) to examine maturational changes in both oscillatory (i.e., periodic) and non-oscillatory (aperiodic, ‘1/*f*-like’) activity in a large cohort of participants ranging from 4-to-12 years of age (N=139, average age=9.41 years, SD=1.95). The EEG signal was parameterised into aperiodic and periodic components, and linear regression models were used to evaluate if chronological age could predict aperiodic exponent and offset, as well as well as peak frequency and power within the alpha and beta ranges. Exponent and offset were found to both decrease with age, while aperiodic-adjusted alpha peak frequency increased with age; however, there was no association between age and peak frequency for the beta band. Age was also unrelated to aperiodic-adjusted spectral power within either the alpha or beta bands, despite both frequency ranges being correlated with the aperiodic signal. Overall, these results highlight the capacity for both periodic and aperiodic features of the EEG to elucidate age-related functional changes within the developing brain.

## 1. Introduction

Electroencephalography (EEG) has proven highly valuable in quantifying neural dynamics and providing critical insights into the physiological processes that underlie key aspects of human cognition and neurodevelopment. Neural oscillations represent a prominent and extensively investigated feature of the EEG record, reflecting synchronised fluctuations in excitability across cortical microcircuits, both within and between broader neuronal networks (Buzsaki & Draguhn, 2004; Cohen, 2017). Decades of research has linked oscillatory activity within the cortex to a broad range of cognitive, perceptual, and developmental processes (Benchenane et al., 2011; Kahana, 2006), while changes in the frequency or amplitude of oscillations can be a sign of pathological neural activity in a number of psychiatric, neurological, and neurodevelopmental disorders (Başar, 2013; Newson & Thiagarajan, 2018; Voytek & Knight, 2015; Wang et al., 2013).

Resting-sate EEG recordings can be used to capture spontaneous, or ‘intrinsic’ activity that occurs in the absence of any overt external stimuli, or task-related neurocognitive processing (Buzsáki et al., 2012; Michel & Murray, 2012). A common trend observed in studies examining maturational changes in resting-state brain rhythms in children and adolescents is a reduction in power with increasing age within lower frequency ranges (i.e., delta and theta bands; Clarke et al., 2001; Gasser et al., 1988; Gómez et al., 2013; John et al., 1980) which is also often also accompanied by a concomitant increase in power within faster rhythms, particularly the alpha and beta bands (Benninger et al., 1984; Gasser et al., 1988; Gómez et al., 2013; Marshall et al., 2002; Saby & Marshall, 2012). In addition to changes in spectral power, the peak frequency of the dominant posterior alpha rhythm also increases with age until around late childhood, or early adulthood (Cellier et al., 2021; Chiang et al., 2011; Eeg-Olofsson et al., 1971; Marshall et al., 2002; Miskovic et al., 2015; Stroganova et al., 1999). These shifts in oscillatory dynamics likely reflect multiple structural and functional neurodevelopmental processes, including differentiation and specialisation of cortical regions/networks, synaptic and axonal pruning, and alterations in excitatory and inhibitory (E/I) circuits (De Bellis et al., 2001; Feinberg & Campbell, 2010; Lujan et al., 2005; Uhlhaas et al., 2010).

The EEG signal, however, reflects not only oscillatory (i.e., periodic) activity, but also additional background aperiodic, or ‘scale-free’ broadband activity, which is present at all frequencies and adheres to a 1/*f* power distribution, whereby spectral power decreases with increasing frequency (Barry & De Blasio, 2021; Donoghue et al., 2020b; He, 2014; Muthukumaraswamy & Liley, 2018; Pritchard, 1992). Despite constituting a large proportion of the spontaneous neural activity recorded from the cortex (Bullock et al., 2003; He et al., 2010), the aperiodic component has until recently received only limited attention in the EEG literature, often being treated as ‘noise’ and regarded as having limited physiological relevance (Donoghue et al., 2020b; He, 2014). Recent work, however, has begun to provide compelling evidence in support of the importance of the aperiodic signal. Studies have shown aperiodic activity to be modulated by task-performance (He et al., 2010), level of arousal (Lendner et al., 2020), and drug-induced states (Colombo et al., 2019; Muthukumaraswamy & Liley, 2018; Waschke et al., 2021). In addition, several studies have shown features of the aperiodic signal to be altered in neurological and psychiatric disease (Molina et al., 2020; Ostlund et al., 2021a; Robertson et al., 2019; Wilkinson & Nelson, 2021).

The aperiodic signal is comprised of two parameters: a spectral slope (henceforth referred to as the aperiodic *exponent*), and an *offset* (Donoghue et al., 2020b). The exponent represents the pattern of power across frequencies, reflecting the steepness of the decay of the power spectrum (Donoghue et al., 2020b), while the offset reflects the broadband shift in power across frequencies (Figure 1B). Emerging research now indicates that these parameters show changes across the lifespan. In adults, a reduction in the exponent (i.e., ‘flatter’ power spectral density [PSD]) with increasing age has been observed across several independent studies (Dave et al., 2018; Merkin et al., 2021; Tran et al., 2020; Voytek et al., 2015). There is also some limited evidence to suggest that these age-dependent changes also occur during childhood. For example, a recent longitudinal EEG study in infants (age ranging between 38 to 203 days) revealed that exponent values decline with age across this early developmental window (Schaworonkow & Voytek, 2021). Additionally, an analysis of magnetoencephalographic (MEG) recordings from a cohort of 24 neurotypical children (mean age = 8.0 years) and 24 adults (mean age = 40.6 years) showed adults to exhibit flatter exponents, and smaller offset values than children (He et al., 2019). A further study containing EEG recordings from both children and young adults (age range 5-21 years), the majority (∼ 88%) of whom had a psychiatric diagnosis, also reported a flattening of the aperiodic exponent, and reduction in offset with increasing age (Tröndle et al., 2020). Similarly, Cellier et al. (2021) recently reported a similar trend in a sample containing both children and adults (age range 3-24 years).

**Figure 1.**
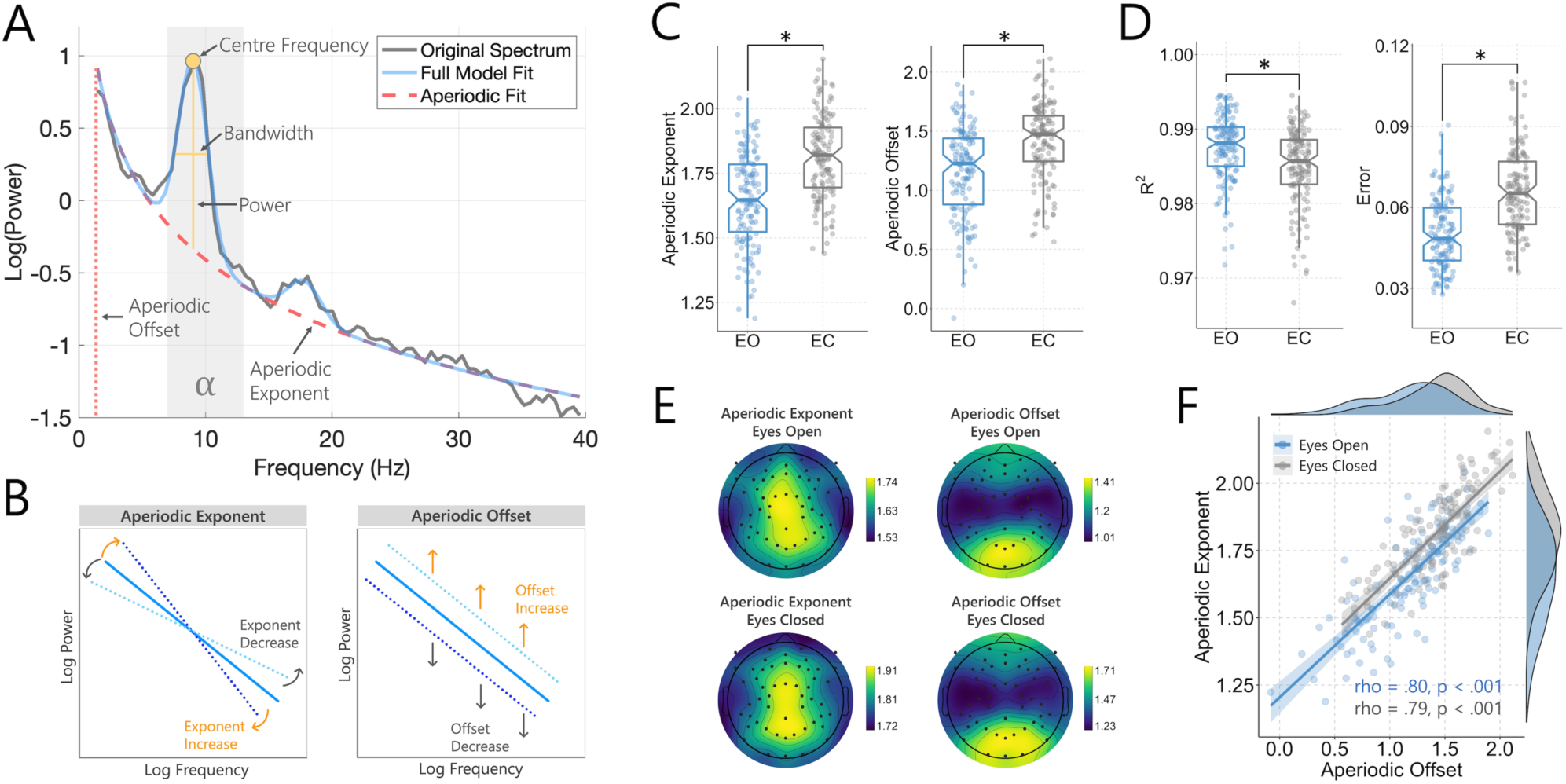
A) Example FOOOF model fit from a single subject showing the aperiodic exponent and offset (marked in blue) across the analysed frequency range (1-40 Hz). The centre frequency, power, and bandwidth are highlighted (arrows) for the oscillatory peak present within the alpha range. B) Graphical illustration demonstrating shifts in the aperiodic exponent and offset. C) Aperiodic exponent and offset values for the eyes open (EO) and eyes closed (EC) recordings. Values are the average across all EEG electrodes. D) R-squared and error values for the model fit for the eyes open and eyes closed recordings (average across all electrodes). E) Topographic plots showing the spatial distribution of mean exponent and offset values across participants. Exponent values were highest near the midline, spanning frontal, central and posterior channels; while offset values were highest over posterior channels. F) Correlation between exponent and offset values (average over all electrodes). There was a strong association between both metrics for the eyes open and eyes closed data.

In sum, neural activity patterns demonstrate various changes across the lifespan. Growing evidence indicates that these alterations do not only reflect shifts in oscillatory dynamics, but also changes within the underlying broadband aperiodic signal. The developmental period spanning early-to-middle childhood is a time of significant and widespread functional and neuroanatomical changes which correspond to vastly increased social and cognitive demands (Bunge & Wright, 2007; Casey et al., 2005). Understanding modifications in both oscillatory and aperiodic neural dynamics within this critical neurodevelopmental period is therefore likely to provide important insight into the physiological processes which take place during this time. The primary aim of the present study was to provide a comprehensive analysis of both periodic and aperiodic components of the spontaneous EEG record in a large cohort of neurotypical children. To achieve this, we employed a recently developed spectral parameterisation approach (Fitting Oscillations and One Over *f* [FOOOF] (Donoghue et al., 2020b)) which enables decomposition of the neural signal into its respective periodic and aperiodic components. This permits narrowband oscillatory dynamics (e.g., power and centre frequency) to be extracted from, and studied independently of, the broadband aperiodic signal. Equally importantly, it further allows explicit measurement of the aperiodic signal, which is likely to be driven by a unique set of neural generators (Donoghue et al., 2021; Ostlund et al., 2021b). Using linear regression models, we examined whether chronological age would predict exponent and offset within the aperiodic signal, as well as the power and centre frequency of the dominant alpha and beta oscillations.

## 2. Methods

### 2.1 Participants

The sample comprised 139 typically developing children (72 male; average age = 9.41 years, SD = 1.95; age range: 4-12 years). All participants were proficient English speakers, and had no history of any neurodevelopmental or neuropsychiatric disorder (as reported by their primary care-giver). Ethical approval was provided by the Deakin University Human Research Ethics Committee (2017-065), while approval to approach public schools was granted by the Victorian Department of Education and Training (2017_003429).

### 2.2 Procedure

Data were collected during a single experimental session conducted either at the university laboratory, or in a quiet room at the participants’ school. Prior to commencement of the study, written consent was obtained from the parent or legal guardian of each child. Details of the experimental protocol were also explained to each child who then agreed to participate. Data reported in this study were collected as part of a larger neurocognitive and electrophysiological investigation into the development of the social brain in early and middle childhood (Bigelow et al., 2021).

### 2.3 EEG data acquisition

EEG data were recorded in a dimly lit room using a 64-channel HydroCel Geodesic Sensor Net (Electrical Geodesics, Inc, USA) containing Ag/AgCl electrodes surrounded by electrolyte-wetted sponges. Data were acquired using NetStation software (version 5.0) via a Net Amps 400 amplifier using a sampling rate of 1 KHz, with data online referenced to the Cz electrode. Prior to the commencement of recording, electrode impedances were checked to ensure they were < 50 KOhms. The resting-state data were recorded for two minutes while participants sat with their eyes open and stared at a fixation cross on a computer screen, and two minutes while participants had their eyes closed.

### 2.4 EEG data analysis

#### 2.4.1 Pre-processing

All pre-processing procedures were performed in Matlab (R2020a; The Mathworks, Massachusetts, USA) incorporating the EEGLAB toolbox (Delorme & Makeig, 2004) along with custom scripts. The raw EEG files were cleaned using the Reduction of Electrophysiological Artifacts (RELAX) pre-processing pipeline (Bailey et al., 2021). This validated and fully automated pipeline uses empirical approaches to identify and reduce artifacts within the data, including the use of both multiple Wiener filters and wavelet enhanced independent component analysis (ICA). Briefly, data were bandpass filtered between 0.5 – 80 Hz (fourth-order Butterworth filter), with a notch filter between 47-53 Hz to remove any line noise, following which any bad channels were removed using a multi-step process including the ‘findNoisyChannels’ function from the PREP pipeline (Bigdely-Shamlo et al., 2015). Data were then subject to multiple Wiener filtering, followed by wavelet-enhanced ICA, with components for cleaning identified using IClabel (Pion-Tonachini et al., 2019). Data were re-referenced to the average of all electrodes ready for further analysis. As a final step, all pre-processed data files were also visually inspected prior to inclusion in the analyses. An overview of the key steps involved in the RELAX pre-processing pipeline can be found in the Supplemental Materials (Figure S1).

#### 2.4.2 Parameterisation of the spectral data

PSD was first calculated separately for each participant and electrode across the continuous EEG using Welch’s method implemented in Matlab (2 second Hamming window, 50% overlap). The FOOOF Python toolbox (version 1.0.0; https://fooof-tools.github.io/fooof/) was then used to parameterize the spectral data through separation of the periodic and aperiodic components of the signal. Using this approach, PSDs are treated as a linear combination of both aperiodic activity and oscillatory peaks with amplitudes that extend above the aperiodic signal (for a detailed overview of this approch see: Donoghue et al., 2020b; Ostlund et al., 2021b). Using a model driven approach, the FOOOF algorithm is able to extract both periodic and aperiodic components within the overall power spectra (Donoghue et al., 2020b). For the present study, we extracted the aperiodic exponent and offset across a broad frequency range between 1 and 40 Hz, similar to prior studies (Cellier et al., 2021; Molina et al., 2020; Ostlund et al., 2021a), and as recommended in the FOOOF documentation in order to allow for reliable estimation of the aperiodic component of the data. Fitting was performed using the ‘fixed’ aperiodic mode due to the absence of a clear ‘knee’ in the power spectrum when the output was visually inspected in log-log space (i.e., the signal was approximately linear across the specified frequency range). Spectral parameterisation settings for the algorithm were: peak width limits = [1, 12], maximum number of peaks = 8, peak threshold = 2, minimum peak height = 0.0. The final FOOOF outputs are the aperiodic exponent and offset values, as well as the centre frequency, power, and bandwidth for the oscillatory component of the signal (see Figure 1A).

#### 2.4.3 Aperiodic Exponent and Offset

The exponent and offset values were extracted from the aperiodic signal for each participant and for each EEG electrode. Prior to statistical analysis, the data were averaged across all scalp electrodes for each participant to generate a ‘global’ exponent and offset value representing the mean signal across the scalp. This approach was chosen as we had no *a priori* hypotheses regarding the scalp distribution of the aperiodic components (Jacob et al., 2021) and also helped to avoid multiple comparisons across electrodes. In instances where significant results were achieved at the global level, we then ran additional analyses across three broad cortical regions using the average signal across electrode clusters covering bilateral anterior (Fp1, Fp2, AFz, AF3, AF4, Fz, F1, F2, F3, F4, F5, F6, F7, F8), central (FCz, FC1, FC2, FC3, FC4, C1, C2, C3, C4, C5, C6, CP1, CP2), and posterior (Pz, P1, P2, P3, P4, P5, P6, P7, P8, POz, PO3, PO4, Oz, O1, O2) channels^*^ (see Figure 2C for a depiction of the EEG cap with the three electrode clusters highlighted).

**Figure 2:**
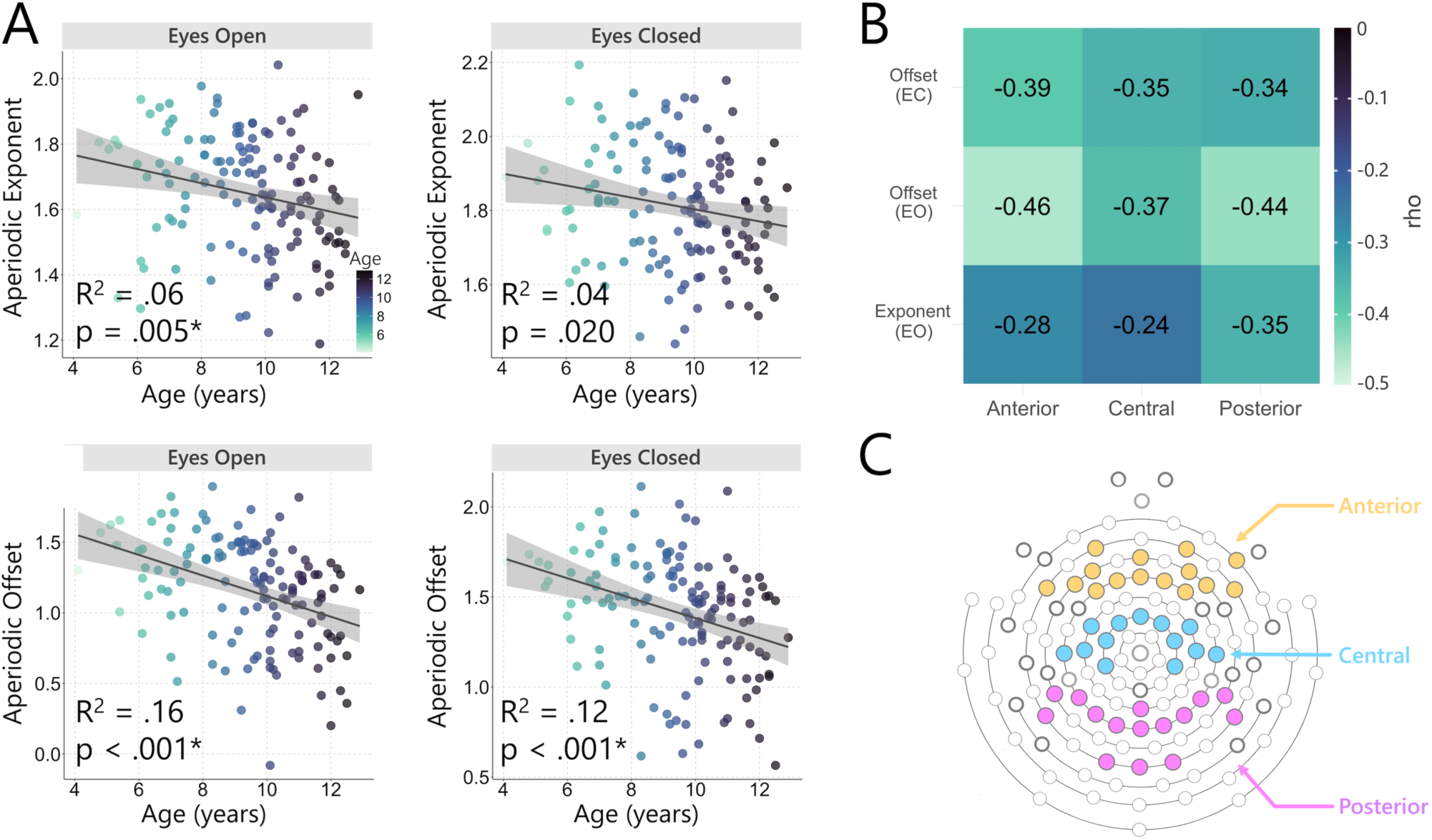
Association between age and aperiodic activity. A) Scatterplot of the aperiodic exponent (upper panel) and offset (lower panel) in relation to age for the eyes open and eyes closed EEG recordings. R-squared (R^2^) and significance values from the regression analyses are shown (asterisk indicates significance after Bonferroni correction). B) Correlations between aperiodic activity and age for each of the anterior, central, and posterior electrode clusters. Correlations reached significance across all three locations. C) EEG electrode cap highlighting the electrodes forming each of the electrode clusters used for the correlations (anterior = yellow, central = blue, posterior = magenta).

#### 2.4.4 Spectral power and centre frequency

Following spectral parameterisation, the *power* and *centre frequency* peak parameters were extracted from the periodic signal for both the alpha (7-13 Hz) and beta (13-30 Hz) frequency ranges. These two frequency ranges were selected based on visual inspection of the power spectra, which indicated clear peaks (i.e., ‘bumps’ in the power spectra) over-and-above the 1/*f*-like decay for most participants (e.g., Figure 1A). Conversely, far fewer participants demonstrated clearly discernible peaks within the canonical delta (1-3 Hz), theta (3-7 Hz), and gamma (>30 Hz) ranges, consistent with other findings (Ostlund et al., 2021b). For statistical analysis, spectral power and centre frequency values were extracted from the midline electrode exhibiting the highest power value (alpha = POz, beta = FCz). Single electrodes, as opposed to electrode clusters, were used in this instance, as the precise number of peaks detected for each electrode differed across subjects, thus prohibiting averaging across a larger ROI. Following removal of the aperiodic components of the signal, an alpha peak was detected in 131 (94.2%) and 138 (99.3%) of participants in the eyes open and eyes closed recordings, respectively; while in the beta range, a peak was successfully detected in 136 (97.8%) and 132 (95.0%) participants for the eyes open and eyes closed conditions, respectively.

### 2.5 Statistical analysis

Statistical analyses were conducted in R (version 4.0.3; R Core Team, 2020). Ordinary least squares regression models were used to predict each of the aperiodic (exponent, offset) and periodic (centre frequency, spectral power) EEG components from chronological age. Regressions were run separately for the eyes open and eyes closed EEG recordings, and for each outcome variable. Residuals diagnostics were performed for all models to assess assumptions (‘olsrr’ package). This included visual inspection of residual Q-Q plots, residual versus fitted values plots, and histograms, as well as Kolmogorov-Smirnov normality tests. In cases of severe violations, the outcome variable was transformed using Yeo-Johnson power transformations (Yeo & Johnson, 2000). As the aperiodic data used the average signal across all electrodes, we also ran further Spearman rank-order correlations to assess for associations between age and exponent and offset values separately across anterior, central, and posterior electrode clusters in instances where regression models were significant. Finally, we ran exploratory correlations to assess for any associations between the aperiodic exponent and offset values, and aperiodic-adjusted spectral power. For all analyses, Bonferroni corrections were used to control for multiple comparisons. For regression models using the aperiodic data, we corrected for four comparisons (2 aperiodic parameters [exponent, offset] x 2 recording conditions [eyes open, eyes closed]; adjusted alpha = .0125). For the periodic data, we corrected for eight comparisons (2 periodic parameters [centre frequency, spectral power] x2 frequencies [alpha, beta] x 2 recording conditions [eyes open, eyes closed]; adjusted alpha = .006). Correlations comparing age and aperiodic activity across specific scalp locations were corrected to account for electrode cluster (3 [anterior, central, posterior] x 2 recording conditions [eyes open, eyes closed]; adjusted alpha = .008). Correlations between exponent and offset were corrected for two correlations (eyes open, eyes closed; adjusted alpha = .025); while correlations between aperiodic activity and spectral power were corrected for eight comparisons (2 periodic parameters [centre frequency, spectral power] x 2 periodic parameters [peak frequency, power], and x 2 recording conditions [eyes open, eyes closed]; adjusted alpha = .006).

## 3. Results

### 3.1 Algorithm performance and characteristics of the aperiodic exponent and offset

The performance of the FOOOF algorithm was assessed via the ‘goodness of fit’ measures, *R*^*2*^ and *Error*, which represent the explained variance, and total error of the model fit, respectively (Donoghue et al., 2020b; Ostlund et al., 2021b). Good model fits for the FOOOF algorithm were observed for both the eyes open (R^2^ = .99, Error = .05) and eyes closed (R^2^ = .98, Error = .07) data (average over all participants/electrodes; Figure 1D). When comparing the eyes open and eyes closed data, R^2^ values were found to be higher for the eyes open, compared to the eyes closed, condition, t(138) = 6.58, p < .001, while Error values were lower for the eyes open, compared to the eyes closed, condition, t(138) = -14.11, p < .001. Topographic plots of the aperiodic exponent and offset (average across all subjects) revealed similar patterns for both the eyes open and eyes closed recordings. Specifically, exponent values showed a relatively widespread distribution, with maximal signal close to the vertex, while offset values were largest across posterior regions of the cortex (Figure 1E). Comparisons between the eyes open and eyes closed recordings also indicated that exponent values were significantly larger (i.e., steeper aperiodic slope) in the eyes closed, compared to the eyes open recordings, t(138) = -12.68, p <.001, with offset values also significantly larger for the eyes closed, compared to the eyes open recordings, t(138) = -14.262, p <.001 (Figure 1C). Finally, exponent and offset values were found to be strongly positively correlated in both the eyes open (rho = .80, p < .001) and eyes closed (rho = .79, p < .001) conditions (Figure 1F).

### 3.1 Association between age and aperiodic activity

Regression models revealed that age predicted the aperiodic exponent for the eyes open EEG recordings, F(1,37) = 8.12, p = .005, R^2^ = .06; however, the eyes closed recordings failed to reach significance after multiple comparison correction, F(1,37) = 5.56, p = .020, R^2^ = .04 (Figure 2A). Age also significantly predicted offset for both the eyes open, F(1,37) = 25.35, p < .001, R^2^ = .16, and eyes closed, F(1,37) = 18.07, p < .001, R^2^ = .12, EEG recordings (Figure 2A). For conditions that reached significance in the regression models, we further examined the relationship between age and aperiodic activity, through correlations using exponent and offset values taken from the average across anterior, central, and posterior scalp locations (see Figure 2C for a depiction of the electrode clusters used). Correlations were significant across each of these three locations (all p < .008), with the strongest association between age and exponent identified posteriorly (*rho* = -.35), and the strongest association between age and offset over the anterior region for both the eyes open (*rho* = -.46) and eyes closed (*rho* = -.39) conditions. Correlation coefficients for all associations are provided in Figure 2B. Additional scatterplots can be found in the Supplementary Materials (Figure S2).

### 3.2 Age-related differences in aperiodic-adjusted centre frequency

Age was found to predict alpha centre frequency across both the eyes open, F(1,129) = 20.73, p <.001, R^2^ = .14, and eyes closed, F(1,136) = 22.31, p <.001, R^2^ = .14, recordings. Specifically, these findings indicate that alpha centre frequency increased systematically with age. In contrast, age did not predict beta centre frequency for either eyes open, F(1,134) = .13, p = .72, R^2^ = .00, or eyes closed, F(1,130) = .51, p = .47, R^2^ = .00, conditions. Scatterplots highlighting the association between age and centre frequency within the alpha and beta bands are presented in Figure 3A.

**Figure 3:**
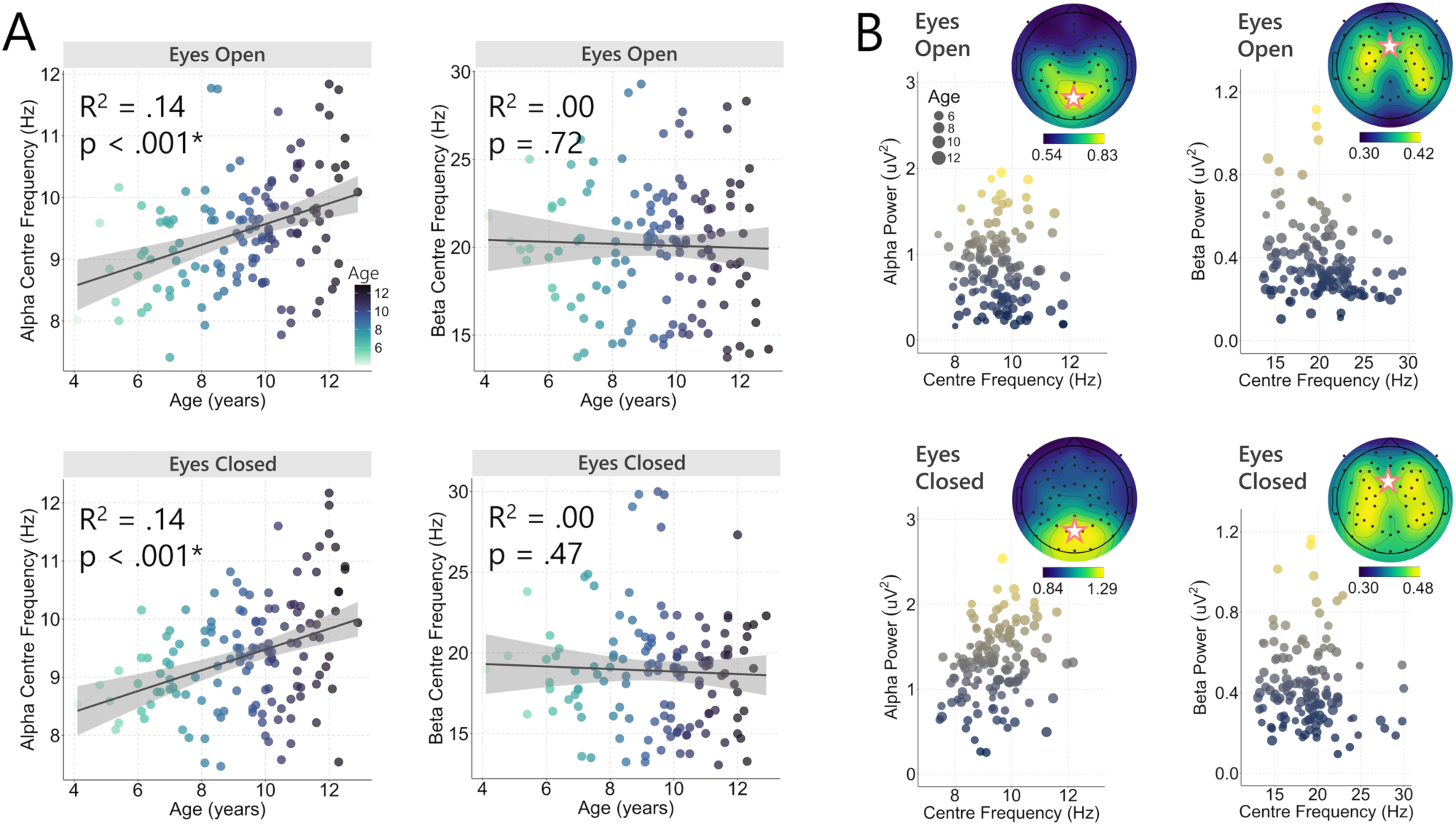
A) Scatter plots of centre frequency for the alpha and beta range in relation to age. Significance values from the regression analyses are shown (asterisk indicates significance after Bonferroni correction). Age was found to significantly predict alpha, but not beta, centre frequency. B) Spectral power plotted in relation to centre frequency for the alpha and beta frequency ranges. Topographic plots show the average power distribution for each of the eyes open and eyes closed recordings for the alpha and beta frequency ranges. Star indicates the electrode used for obtaining the power and centre frequency values used in the analyses (alpha = POz electrode, beta = FCz electrode).

### 3.3 Association between age and aperiodic-adjusted spectral power

We assessed whether age could predict the power of the detected oscillatory peaks in the alpha and beta ranges after removal of the aperiodic signal. No association between age and power was found for the alpha (eyes open: F(1,129) = .08, p = .78, R^2^ = .00; eyes closed: F(1,136) = 3.02, p = .08, R^2^ = .02), or beta (eyes open: F(1,134) = 1.67, p = .20, R^2^ = .01; eyes closed: F(1,130) = 4.33, p = .04, R^2^ = .03) frequencies (for scatterplots, see Supplementary Figure S3). Figure 3B depicts spectral power in relation to centre frequency for the alpha and beta frequencies. Given recent work identifying a potential association the aperiodic signal and aperiodic-adjusted spectral power and peak frequency (He et al., 2019; Merkin et al., 2021), exploratory analyses were also run comparing exponent and offset (average across all electrodes) with power and centre frequency within the alpha and beta ranges (using the midline electrode exhibiting the greatest amplitude [alpha = POz, beta = FCz]). We found significant weak-to-moderate positive correlations between exponent and offset and aperiodic-adjusted spectral power in both the alpha and beta bands for both the eyes open and eyes closed data (Bonferroni corrected; all p < .006; Figure 4). When comparing aperiodic activity with centre frequency, the only significant result was a modest negative association between beta power in the eyes open recordings and exponent (rho = -.24, p = .005) and offset (rho = -.26, p = .002) (Supplementary Figure S4).

**Figure 4:**
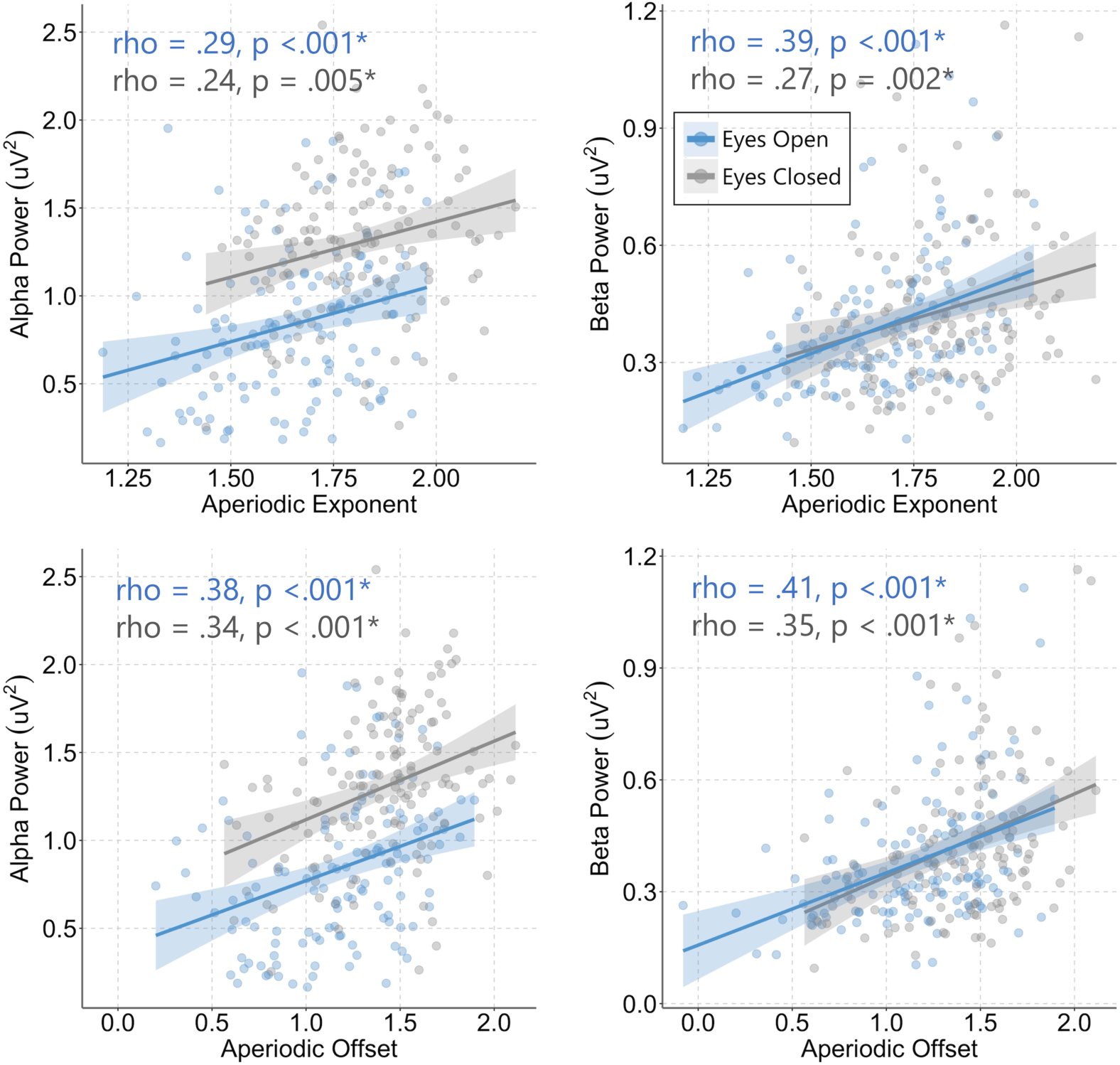
Scatterplots depicting the association between aperiodic activity and aperiodic-adjusted oscillatory power. Both exponent and offset positively correlated with spectral power in both the alpha and beta bands across the eyes open and eyes closed conditions. Asterisks indicate a significant correlation after Bonferroni correction.

## 4. Discussion

The aim of the present study was to characterise neurodevelopmental changes across both the periodic and aperiodic components of the spontaneous EEG signal in early-to-middle childhood. To achieve this, we applied a spectral parameterization approach to disentangle key features (i.e., centre frequency and power) of narrowband oscillations in the alpha and beta band from the broader aperiodic signal. Using regression models, we then examined if participants’ age could predict the aperiodic exponent and offset, as well as power and centre frequency within the alpha and beta frequencies. Several key findings emerged from this investigation. First, we found chronological age to be a predictor of both aperiodic exponent and offset, with older children having smaller exponent values (i.e., flatter 1/*f* spectral slope) and reduced offset values, compared to younger children. Second, we found that age was also able to predict centre frequency of the alpha (but not beta) band, with older children displaying faster peak frequencies. Finally, age was unable to predict spectral power in either the alpha or beta bands; however, power within these bands significantly correlated with both aperiodic exponent and offset.

### 4.1 Age predicts aperiodic properties of the EEG signal

The present results are indicative of an association between children’s chronological age and the aperiodic properties of the EEG signal. Specifically, exponent values, extracted from the eyes open recordings, were shown to decline as a function of age; however, this association failed to reach significance in the eyes closed data after multiple comparison correction. Offset also declined with age, with this result significant for both the eyes open and eyes closed recordings. These findings largely corroborate previous work also indicating age-related changes in the aperiodic EEG signal. An initial study by Voytek et al. (2015) that examined EEG recorded during a visual working memory task found that the slope of the 1/f signal was less negative (i.e., ‘flatter’ aperiodic exponent) in older (60-70 years), compared to younger (20-30 years) participants. Similar results were also reported by Tran et al. (2020) comparing participants across the same age groups as Voytek et al. (2015), but instead using pretrial baseline EEG recorded during a cognitive task. Consistent with Voytek et al. (2015) these authors also found that exponent was flatter in the older, compared to younger, group. These key findings of smaller exponent and reduced offset in older individuals were further recently supported by Merkin et al. (2021) using eyes closed resting-state EEG in younger (18-35 years) and older (50-86 years) adults. Our present findings add to this emerging body of evidence in adult populations by demonstrating reduced exponent and offset with age in a large cohort spanning early-to-middle childhood (4-to-12 years). These findings were present both when using the signal averaged across the entire scalp, and when using electrode clusters covering anterior, central, and posterior regions separately (Figure 2B and 2C), thus suggesting that systematic variations with age are a relatively widespread phenomenon. Importantly, our findings also replicate recent observations of flattening of the aperiodic exponent with age in infancy (Schaworonkow & Voytek, 2021), as well as in cohorts with ages ranging from childhood into adulthood (Cellier et al., 2021; Donoghue et al., 2020a; He et al., 2019; Tröndle et al., 2020). The present results, in conjunction with previous findings, are therefore supportive of quantitative neurodevelopmental changes in the aperiodic component of the EEG signal.

Although precise neurobiological substrate of aperiodic activity remains uncertain, evidence suggests that a flatter exponent reflects increased asynchronous background neuronal firing (i.e., neural ‘noise’) which is theorised to be driven by an increased E/I ratio (Voytek & Knight, 2015; Voytek et al., 2015). This has recently been supported via both *in silico* models (Gao et al., 2017), as well as neural recordings demonstrating modulation of the spectral exponent through administration of pharmacological agents known to either increase inhibition (e.g., propofol), or increase excitation (ketamine) (Gao et al., 2017; Lendner et al., 2020; Waschke et al., 2021). Hence, it is possible that the age-dependent exponent reductions observed here reflect, to some extent, maturational changes in E/I balance occurring throughout this neurodevelopmental period, possibly representing a shift towards increased excitatory tone within neural circuits as children mature. However, further work is needed to elucidate the precise cellular and molecular mechanisms that underlie these neurodevelopmental shifts in aperiodic activity. Future studies could combine analysis of the EEG-derived aperiodic signal with neuroimaging techniques capable of quantifying excitatory and inhibitory neurotransmitter concentrations (i.e., γ-aminobutyric acid [GABA] and glutamate) within the brain, such as magnetic resonance spectroscopy (MRS) (Harris et al., 2017; Thakkar et al., 2017). For instance, there is some limited evidence that GABA levels increase during neurodevelopment (Porges et al., 2021), however, exactly how this finding ties in with markers of neural excitability remains to be established. Multi-modal approaches combining neurostimulation with electrophysiology, such as combined transcranial magnetic stimulation and EEG (TMS-EEG), could also be utilised to probe associations between cortical excitability (via TMS-evoked potentials) and aperiodic activity across specific cortical targets (Hill et al., 2016; Tremblay et al., 2019).

The observation of age-related reductions in offset also warrants further investigation. Intracranial local field potential recordings from patients undergoing neurosurgery provide compelling evidence that broadband power shifts are positively correlated to neuronal population spiking (Manning et al., 2009), with similar findings also observed in macaques (Ray & Maunsell, 2011). Hence, our present observation of a reduction in aperiodic offset with increasing age could be tentatively interpreted to reflect a maturational decline in the spiking rate of cortical neurons. In keeping with the observed changes in aperiodic exponent, this effect appears to be a relatively global phenomenon, given that results were taken from the average of the aperiodic signal across all scalp electrodes. More broadly, these findings also appear consistent with previous observations of reduced broadband power throughout childhood and into adulthood (Gomez et al., 2017; Segalowitz et al., 2010). It is possible that reductions in cortical grey matter volume that occur during childhood, likely the result of maturational ‘synaptic pruning’-like processes (Paolicelli et al., 2011; Paus et al., 2008; Pfefferbaum et al., 1994), are responsible for these findings. However, we note that cautious interpretation is warranted, as changes in skull conductivity with age might also contribute to these observations when using EEG recordings (Gomez et al., 2017; Hoekema et al., 2003). In any event, the results of He et al. (2019), which also reported an age-related decline in offset, lend support to such changes being genuine neural phenomena, given that these authors used MEG recordings, which are largely unaffected by the electrical resistivity of the skull (Wolters et al., 2006).

### 4.2 Age predicts alpha centre frequency, but not power

Participants’ age was able to predict the aperiodic-adjusted alpha frequency, with older children showing increased alpha peak frequency. This finding aligns with previously documented observations of alpha frequency with age both across childhood (Dickinson et al., 2018; Eeg-Olofsson et al., 1971; Marshall et al., 2002; Miskovic et al., 2015; Somsen et al., 1997), and into adolescence and early adulthood (Chiang et al., 2011; Cragg et al., 2011). The posterior alpha rhythm first manifests on the EEG record at around 3 months of age, with a peak frequency between 3-5 Hz, which increases to 6-7 Hz by one year of age (Saby & Marshall, 2012), and continues to increase throughout childhood until reaching a peak between 8-12 Hz in early adulthood, after which it steadily declines with age (Chiang et al., 2011; Hashemi et al., 2016). It has been theorised that the increase in alpha frequency seen throughout childhood might represent an increase in the speed at which interconnected neural populations are able to communicate, as a result of greater myelination and axon diameter (Segalowitz et al., 2010; Thorpe et al., 2016). A relationship between increasing alpha frequency and the development of large-scale oscillatory networks would also be in alignment with studies that have shown associations between peak alpha frequency and cognitive function in children (Carter Leno et al., 2021; Dickinson et al., 2018). Our present findings also corroborate recent reports of age-related changes in aperiodic-adjusted alpha centre frequency. Specifically, Cellier et al. (2021) showed that peak frequency within the 4-12 Hz range increased with age in a cohort which included both children and adults (3-24 years of age); while He et al. (2019) reported a positive association between age and alpha centre frequency in a small sample (N = 24) of children using MEG recordings. The same trend was also reported by Carter-Leno et al. (2021) in a longitudinal sample of young children from 1 to 3 years of age. Our findings extend these observations by demonstrating age-related shifts in alpha centre frequency using both eyes open and eyes closed data from a large (N = 139) cohort of children spanning early-to-middle childhood. Our results were also specific to the alpha band, with no evidence of an association between age and beta centre frequency. The maturational changes observed in alpha frequency, however, did not extend to spectral power in either the alpha or beta frequencies. Although a number of studies have shown associations between power in various canonical frequencies and age, considerable heterogeneity exists, with results likely to be strongly contingent on the specific age range investigated (for review see: Segalowitz et al., 2010). Importantly, the majority of past research examining narrow-band oscillatory power has failed to account for the potential influence of the aperiodic signal, which risks conflating these two separate phenomena (Donoghue et al., 2020a; Donoghue et al., 2021). Recent work examining peak alpha frequency in adulthood also found no age-related changes after accounting for the aperiodic signal (Merkin et al., 2021).

### 4.3 Limitations and future directions

The present study has some limitations. First, as we used resting-state EEG, the present results are limited to spontaneous neural activity. Whilst this is valuable for understanding intrinsic (i.e., stimulus free) dynamics, future work could further extend these findings using task-related paradigms. This might be particularly useful to help identify relationships between periodic and aperiodic neural dynamics and specific neurocognitive processes. For example, recent work has identified aperiodic activity as a predictor of working memory performance (Donoghue et al., 2020b) and cognitive processing speed (Ouyang et al., 2020). Second, we parameterised our data between 1-40 Hz. We chose this range to reduce the presence of non-neural artefacts (e.g., electromyographic activity, or microsaccades) which often occur at higher frequencies (Goncharova et al., 2003; Muthukumaraswamy, 2013; Yuval-Greenberg et al., 2008). While this still represents a broad frequency range, and is consistent with other EEG studies utilising spectral parameterisation (Carter Leno et al., 2021; Cellier et al., 2021; Merkin et al., 2021; Robertson et al., 2019), future work could extend these analyses to even wider frequency ranges to capture higher frequency activity (e.g., > 40 Hz). This might be particularly useful for more directly comparing MEG and EEG derived data with results from local field potential and electrocorticography recordings (e.g., Gao et al., 2017; Halgren et al., 2021). Finally, emerging evidence indicating that the aperiodic exponent might act as a non-invasive measure of E/I balance (Gao et al., 2017; Waschke et al., 2021) opens exciting possibilities for research into the neurobiology of developmental and neuropsychiatric disorders linked to dysfunction within excitatory and inhibitory circuits, such as autism and schizophrenia (Foss-Feig et al., 2017).

## 4.4 Conclusion

The present results highlight several key maturational effects on the spontaneous EEG in recorded in a large sample of participants spanning early-to-middle childhood (4-to-12 years). Across this age-range, both aperiodic exponent and offset were shown to decrease with age. Further, aperiodic-adjusted peak alpha frequency increased with age, while no effect of age was observed for the beta band. Finally, age was not shown to predict either aperiodic-adjusted alpha, or beta power. These results provide support for nuanced approaches aiming to examine neural dynamics within neurodevelopmental cohorts, which disentangle narrow-band oscillatory features from broadband aperiodic activity.

## Abbreviations

EEG: electroencephalography
E/I: excitation/inhibition
FOOOF: fitting oscillations and one over f
GABA: γ-aminobutyric acid
ICA: independent component analysis
MEG: magnetoencephalography
PSD: power spectral density
ROI: region of interest
TMS: transcranial magnetic stimulation.

## Footnotes

* Note: International 10-10 electrode positions are stated for ease of interpretation. Channels listed are those which best approximate the sensor positions used by the Geodesic Sensor Net.

## REFERENCES

Bailey, N. W., Biabani, M., Hill, A. T., Rogasch, N. C., McQueen, B., & Fitzgerald, P. B. (2021). Introducing RELAX (the Reduction of Electrophysiological Artifacts): A fully automatic pre-processing pipeline for EEG data. In Preparation.

Barry, R. J., & De Blasio, F. M. (2021). Characterizing pink and white noise in the human electroencephalogram. J Neural Eng. doi:10.1088/1741-2552/abe399

Başar, E. (2013). Brain oscillations in neuropsychiatric disease. Dialogues in clinical neuroscience, 15(3), 291–300. doi:10.31887/DCNS.2013.15.3/ebasar

Benchenane, K., Tiesinga, P. H., & Battaglia, F. P. (2011). Oscillations in the prefrontal cortex: a gateway to memory and attention. Curr Opin Neurobiol, 21(3), 475–485. doi:10.1016/j.conb.2011.01.004

Benninger, C., Matthis, P., & Scheffner, D. (1984). EEG development of healthy boys and girls. Results of a longitudinal study. Electroencephalography and Clinical Neurophysiology, 57(1), 1–12. doi:https://doi.org/10.1016/0013-4694(84)90002-6

Bigdely-Shamlo, N., Mullen, T., Kothe, C., Su, K. M., & Robbins, K. A. (2015). The PREP pipeline: standardized preprocessing for large-scale EEG analysis. Front Neuroinform, 9, 16. doi:10.3389/fninf.2015.00016

Bullock, T. H., McClune, M. C., & Enright, J. T. (2003). Are the electroencephalograms mainly rhythmic? Assessment of periodicity in wide-band time series. Neuroscience, 121(1), 233–252. doi:https://doi.org/10.1016/S0306-4522(03)00208-2

Bunge, S. A., & Wright, S. B. (2007). Neurodevelopmental changes in working memory and cognitive control. Curr Opin Neurobiol, 17(2), 243–250. doi:10.1016/j.conb.2007.02.005

Buzsáki, G., Anastassiou, C. A., & Koch, C. (2012). The origin of extracellular fields and currents — EEG, ECoG, LFP and spikes. Nature Reviews Neuroscience, 13(6), 407–420. doi:http://www.nature.com/nrn/journal/v13/n6/suppinfo/nrn3241_S1.html

Buzsaki, G., & Draguhn, A. (2004). Neuronal oscillations in cortical networks. Science, 304(5679), 1926–1929. doi:10.1126/science.1099745

Carter Leno, V., Pickles, A., van Noordt, S., Huberty, S., Desjardins, J., Webb, S. J., … Team, B. (2021). 12-Month peak alpha frequency is a correlate but not a longitudinal predictor of non-verbal cognitive abilities in infants at low and high risk for autism spectrum disorder. Dev Cogn Neurosci, 48, 100938. doi:10.1016/j.dcn.2021.100938

Casey, B. J., Tottenham, N., Liston, C., & Durston, S. (2005). Imaging the developing brain: what have we learned about cognitive development? Trends Cogn Sci, 9(3), 104–110. doi:10.1016/j.tics.2005.01.011

Cellier, D., Riddle, J., Petersen, I., & Hwang, K. (2021). The development of theta and alpha neural oscillations from ages 3 to 24 years. Dev Cogn Neurosci. doi:10.1016/j.dcn.2021.100969

Chiang, A. K., Rennie, C. J., Robinson, P. A., van Albada, S. J., & Kerr, C. C. (2011). Age trends and sex differences of alpha rhythms including split alpha peaks. Clin Neurophysiol, 122(8), 1505–1517. doi:10.1016/j.clinph.2011.01.040

Clarke, A. R., Barry, R. J., McCarthy, R., & Selikowitz, M. (2001). Age and sex effects in the EEG: development of the normal child. Clinical Neurophysiology, 112(5), 806–814. doi:https://doi.org/10.1016/S1388-2457(01)00488-6

Cohen, M. X. (2017). Where Does EEG Come From and What Does It Mean? Trends Neurosci, 40(4), 208–218. doi:10.1016/j.tins.2017.02.004

Colombo, M. A., Napolitani, M., Boly, M., Gosseries, O., Casarotto, S., Rosanova, M., … Sarasso, S. (2019). The spectral exponent of the resting EEG indexes the presence of consciousness during unresponsiveness induced by propofol, xenon, and ketamine. Neuroimage, 189, 631–644. doi:10.1016/j.neuroimage.2019.01.024

Cragg, L., Kovacevic, N., McIntosh, A. R., Poulsen, C., Martinu, K., Leonard, G., & Paus, T. (2011). Maturation of EEG power spectra in early adolescence: a longitudinal study. Developmental Science, 14(5), 935–943. doi:https://doi.org/10.1111/j.1467-7687.2010.01031.x

Dave, S., Brothers, T. A., & Swaab, T. Y. (2018). 1/f neural noise and electrophysiological indices of contextual prediction in aging. Brain Res, 1691, 34–43. doi:10.1016/j.brainres.2018.04.007

De Bellis, M. D., Keshavan, M. S., Beers, S. R., Hall, J., Frustaci, K., Masalehdan, A., … Boring, A. M. (2001). Sex Differences in Brain Maturation during Childhood and Adolescence. Cerebral Cortex, 11(6), 552–557. doi:10.1093/cercor/11.6.552

Delorme, A., & Makeig, S. (2004). EEGLAB: an open source toolbox for analysis of single-trial EEG dynamics including independent component analysis. J Neurosci Methods, 134(1), 9–21. doi:10.1016/j.jneumeth.2003.10.009

Dickinson, A., DiStefano, C., Senturk, D., & Jeste, S. S. (2018). Peak alpha frequency is a neural marker of cognitive function across the autism spectrum. Eur J Neurosci, 47(6), 643–651. doi:10.1111/ejn.13645

Donoghue, T., Dominguez, J., & Voytek, B. (2020a). Electrophysiological Frequency Band Ratio Measures Conflate Periodic and Aperiodic Neural Activity. eNeuro, 7(6). doi:10.1523/ENEURO.0192-20.2020

Donoghue, T., Haller, M., Peterson, E. J., Varma, P., Sebastian, P., Gao, R., … Voytek, B. (2020b). Parameterizing neural power spectra into periodic and aperiodic components. Nat Neurosci, 23(12), 1655–1665. doi:10.1038/s41593-020-00744-x

Donoghue, T., Schaworonkow, N., & Voytek, B. (2021). Methodological considerations for studying neural oscillations. Eur J Neurosci. doi:10.1111/ejn.15361

Eeg-Olofsson, O., Petersén, I., & Selldén, U. (1971). The development of the electroencephalogram in normal children from the age of 1 through 15 years. Paroxysmal activity. Neuropadiatrie, 2(4), 375–404. doi:10.1055/s-0028-1091791

Feinberg, I., & Campbell, I. G. (2010). Sleep EEG changes during adolescence: an index of a fundamental brain reorganization. Brain Cogn, 72(1), 56–65. doi:10.1016/j.bandc.2009.09.008

Foss-Feig, J. H., Adkinson, B. D., Ji, J. L., Yang, G., Srihari, V. H., McPartland, J. C., … Anticevic, A. (2017). Searching for Cross-Diagnostic Convergence: Neural Mechanisms Governing Excitation and Inhibition Balance in Schizophrenia and Autism Spectrum Disorders. Biol Psychiatry, 81(10), 848–861. doi:10.1016/j.biopsych.2017.03.005

Gao, R., Peterson, E. J., & Voytek, B. (2017). Inferring synaptic excitation/inhibition balance from field potentials. Neuroimage, 158, 70–78. doi:10.1016/j.neuroimage.2017.06.078

Gasser, T., Verleger, R., Bächer, P., & Sroka, L. (1988). Development of the EEG of school-age children and adolescents. I. Analysis of band power. Electroencephalography and Clinical Neurophysiology, 69(2), 91–99. doi:https://doi.org/10.1016/0013-4694(88)90204-0

Gómez, C. M, Pérez-Macías, J., Poza, J., Fernández, A., & Hornero, R. (2013). Spectral changes in spontaneous MEG activity across the lifespan. J Neural Eng, 10(6), 066006. doi:10.1088/1741-2560/10/6/066006

Gomez, C. M., Rodriguez-Martinez, E. I., Fernandez, A., Maestu, F., Poza, J., & Gomez, C. (2017). Absolute Power Spectral Density Changes in the Magnetoencephalographic Activity During the Transition from Childhood to Adulthood. Brain Topogr, 30(1), 87–97. doi:10.1007/s10548-016-0532-0

Goncharova, I. I., McFarland, D. J., Vaughan, T. M., & Wolpaw, J. R. (2003). EMG contamination of EEG: spectral and topographical characteristics. Clinical Neurophysiology, 114(9), 1580–1593. doi:https://doi.org/10.1016/S1388-2457(03)00093-2

Halgren, M., Kang, R., Voytek, B., Ulbert, I., Fabo, D., Eross, L. G., … Cash, S. S. (2021). The timescale and magnitude of aperiodic activity decreases with cortical depth in humans, macaques and mice. bioRxiv, 2021.2007.2028.454235. doi:10.1101/2021.07.28.454235

Harris, A. D., Saleh, M. G., & Edden, R. A. (2017). Edited (1) H magnetic resonance spectroscopy in vivo: Methods and metabolites. Magn Reson Med, 77(4), 1377–1389. doi:10.1002/mrm.26619

Hashemi, A., Pino, L. J., Moffat, G., Mathewson, K. J., Aimone, C., Bennett, P. J., … Sekuler, A. B. (2016). Characterizing Population EEG Dynamics throughout Adulthood. eNeuro, 3(6). doi:10.1523/ENEURO.0275-16.2016

He, B. J. (2014). Scale-free brain activity: past, present, and future. Trends Cogn Sci, 18(9), 480–487. doi:10.1016/j.tics.2014.04.003

He, B. J., Zempel, J. M., Snyder, A. Z., & Raichle, M. E. (2010). The temporal structures and functional significance of scale-free brain activity. Neuron, 66(3), 353–369. doi:10.1016/j.neuron.2010.04.020

He, W., Donoghue, T., Sowman, P. F., Seymour, R. A., Brock, J., Crain, S., … Hillebrand, A. (2019). Co-Increasing Neuronal Noise and Beta Power in the Developing Brain. doi:10.1101/839258

Hill, A. T., Rogasch, N. C., Fitzgerald, P. B., & Hoy, K. E. (2016). TMS-EEG: A window into the neurophysiological effects of transcranial electrical stimulation in non-motor brain regions. Neurosci Biobehav Rev, 64, 175–184. doi:10.1016/j.neubiorev.2016.03.006

Hoekema, R., Wieneke, G. H., Leijten, F. S. S., van Veelen, C. W. M., van Rijen, P. C., Huiskamp, G. J. M., … van Huffelen, A. C. (2003). Measurement of the Conductivity of Skull, Temporarily Removed During Epilepsy Surgery. Brain Topogr, 16(1), 29–38. doi:10.1023/A:1025606415858

Jacob, M. S., Roach, B. J., Sargent, K., Mathalon, D. H., & Ford, J. M. (2021). Aperiodic measures of neural excitability are associated with anticorrelated hemodynamic networks at rest: a combined EEG-fMRI study. doi:10.1101/2021.01.30.427861

John, E., Ahn, H., Prichep, L., Trepetin, M., Brown, D., & Kaye, H. (1980). Developmental equations for the electroencephalogram. Science, 210(4475), 1255–1258. doi:10.1126/science.7434026

Kahana, M. J. (2006). The cognitive correlates of human brain oscillations. J Neurosci, 26(6), 1669–1672. doi:10.1523/JNEUROSCI.3737-05c.2006

Lendner, J. D., Helfrich, R. F., Mander, B. A., Romundstad, L., Lin, J. J., Walker, M. P., … Knight, R. T. (2020). An electrophysiological marker of arousal level in humans. Elife, 9. doi:10.7554/eLife.55092

Lujan, R., Shigemoto, R., & Lopez-Bendito, G. (2005). Glutamate and GABA receptor signalling in the developing brain. Neuroscience, 130(3), 567–580. doi:10.1016/j.neuroscience.2004.09.042

Manning, J. R., Jacobs, J., Fried, I., & Kahana, M. J. (2009). Broadband shifts in local field potential power spectra are correlated with single-neuron spiking in humans. J Neurosci, 29(43), 13613–13620. doi:10.1523/JNEUROSCI.2041-09.2009

Marshall, P. J., Bar-Haim, Y., & Fox, N. A. (2002). Development of the EEG from 5 months to 4 years of age. Clinical Neurophysiology, 113(8), 1199–1208. doi:https://doi.org/10.1016/S1388-2457(02)00163-3

Merkin, A., Sghirripa, S., Graetz, L., Smith, A. E., Hordacre, B., Harris, R., … Goldsworthy, M. (2021). Age differences in aperiodic neural activity measured with resting EEG. bioRxiv, 2021.2008.2031.458328. doi:10.1101/2021.08.31.458328

Michel, C. M., & Murray, M. M. (2012). Towards the utilization of EEG as a brain imaging tool. Neuroimage, 61(2), 371–385. doi:10.1016/j.neuroimage.2011.12.039

Miskovic, V., Ma, X., Chou, C. A., Fan, M., Owens, M., Sayama, H., & Gibb, B. E. (2015). Developmental changes in spontaneous electrocortical activity and network organization from early to late childhood. Neuroimage, 118, 237–247. doi:10.1016/j.neuroimage.2015.06.013

Molina, J. L., Voytek, B., Thomas, M. L., Joshi, Y. B., Bhakta, S. G., Talledo, J. A., … Light, G. A. (2020). Memantine Effects on Electroencephalographic Measures of Putative Excitatory/Inhibitory Balance in Schizophrenia. Biological Psychiatry: Cognitive Neuroscience and Neuroimaging, 5(6), 562–568. doi:10.1016/j.bpsc.2020.02.004

Muthukumaraswamy, S. D. (2013). High-frequency brain activity and muscle artifacts in MEG/EEG: a review and recommendations. Front Hum Neurosci, 7, 138. doi:10.3389/fnhum.2013.00138

Muthukumaraswamy, S. D., & Liley, D. T. (2018). 1/f electrophysiological spectra in resting and drug-induced states can be explained by the dynamics of multiple oscillatory relaxation processes. Neuroimage, 179, 582–595. doi:10.1016/j.neuroimage.2018.06.068

Newson, J. J., & Thiagarajan, T. C. (2018). EEG Frequency Bands in Psychiatric Disorders: A Review of Resting State Studies. Front Hum Neurosci, 12, 521. doi:10.3389/fnhum.2018.00521

Ostlund, B. D., Alperin, B. R., Drew, T., & Karalunas, S. L. (2021a). Behavioral and cognitive correlates of the aperiodic (1/f-like) exponent of the EEG power spectrum in adolescents with and without ADHD. Dev Cogn Neurosci, 48, 100931. doi:10.1016/j.dcn.2021.100931

Ostlund, B. D., Donoghue, T., Anaya, B., Gunther, K. E., Karalunas, S. L., Voytek, B., & Perez-Edgar, K. E. (2021b). Spectral parameterization for studying neurodevelopment: How and why. PsyArXiv. doi:https://doi.org/10.31234/osf.io/btqyk

Ouyang, G., Hildebrandt, A., Schmitz, F., & Herrmann, C. S. (2020). Decomposing alpha and 1/f brain activities reveals their differential associations with cognitive processing speed. Neuroimage, 205, 116304. doi:10.1016/j.neuroimage.2019.116304

Paolicelli, R. C., Bolasco, G., Pagani, F., Maggi, L., Scianni, M., Panzanelli, P., … Gross, C. T. (2011). Synaptic pruning by microglia is necessary for normal brain development. Science, 333(6048), 1456–1458. doi:10.1126/science.1202529

Paus, T., Keshavan, M., & Giedd, J. N. (2008). Why do many psychiatric disorders emerge during adolescence? Nature Reviews Neuroscience, 9(12), 947–957. doi:10.1038/nrn2513

Pfefferbaum, A., Mathalon, D. H., Sullivan, E. V., Rawles, J. M., Zipursky, R. B., & Lim, K. O. (1994). A Quantitative Magnetic Resonance Imaging Study of Changes in Brain Morphology From Infancy to Late Adulthood. Arch Neurol, 51(9), 874–887. doi:10.1001/archneur.1994.00540210046012

Pion-Tonachini, L., Kreutz-Delgado, K., & Makeig, S. (2019). ICLabel: An automated electroencephalographic independent component classifier, dataset, and website. Neuroimage, 198, 181–197. doi:10.1016/j.neuroimage.2019.05.026

Porges, E. C., Jensen, G., Foster, B., Edden, R. A., & Puts, N. A. (2021). The trajectory of cortical GABA across the lifespan, an individual participant data meta-analysis of edited MRS studies. Elife, 10. doi:10.7554/eLife.62575

Pritchard, W. S. (1992). The Brain in Fractal Time: 1/F-Like Power Spectrum Scaling of the Human Electroencephalogram. International Journal of Neuroscience, 66(1-2), 119-129. doi:10.3109/00207459208999796

R Core Team. (2020). R: A Language Environment for Statistical Computing. Vienna, Austria: R Foundation for Statistical Computing.

Ray, S., & Maunsell, J. H. (2011). Different origins of gamma rhythm and high-gamma activity in macaque visual cortex. PLoS Biol, 9(4), e1000610. doi:10.1371/journal.pbio.1000610

Robertson, M. M., Furlong, S., Voytek, B., Donoghue, T., Boettiger, C. A., & Sheridan, M. A. (2019). EEG power spectral slope differs by ADHD status and stimulant medication exposure in early childhood. J Neurophysiol, 122(6), 2427–2437. doi:10.1152/jn.00388.2019

Saby, J. N., & Marshall, P. J. (2012). The Utility of EEG Band Power Analysis in the Study of Infancy and Early Childhood. Developmental Neuropsychology, 37(3), 253–273. doi:10.1080/87565641.2011.614663

Schaworonkow, N., & Voytek, B. (2021). Longitudinal changes in aperiodic and periodic activity in electrophysiological recordings in the first seven months of life. Dev Cogn Neurosci, 47, 100895. doi:10.1016/j.dcn.2020.100895

Segalowitz, S. J., Santesso, D. L., & Jetha, M. K. (2010). Electrophysiological changes during adolescence: a review. Brain Cogn, 72(1), 86–100. doi:10.1016/j.bandc.2009.10.003

Somsen, R. J. M., van’t Klooster, B. J., van der Molen, M. W., van Leeuwen, H. M. P., & Licht, R. (1997). Growth spurts in brain maturation during middle childhood as indexed by EEG power spectra. Biol Psychol, 44(3), 187–209. doi:https://doi.org/10.1016/S0301-0511(96)05218-0

Stroganova, T. A., Orekhova, E. V., & Posikera, I. N. (1999). EEG alpha rhythm in infants. Clinical Neurophysiology, 110(6), 997–1012. doi:https://doi.org/10.1016/S1388-2457(98)00009-1

Thakkar, K. N., Rosler, L., Wijnen, J. P., Boer, V. O., Klomp, D. W., Cahn, W., … Neggers, S. F. (2017). 7T Proton Magnetic Resonance Spectroscopy of Gamma-Aminobutyric Acid, Glutamate, and Glutamine Reveals Altered Concentrations in Patients With Schizophrenia and Healthy Siblings. Biol Psychiatry, 81(6), 525–535. doi:10.1016/j.biopsych.2016.04.007

Thorpe, S. G., Cannon, E. N., & Fox, N. A. (2016). Spectral and source structural development of mu and alpha rhythms from infancy through adulthood. Clin Neurophysiol, 127(1), 254–269. doi:10.1016/j.clinph.2015.03.004

Tran, T. T., Rolle, C. E., Gazzaley, A., & Voytek, B. (2020). Linked Sources of Neural Noise Contribute to Age-related Cognitive Decline. J Cogn Neurosci, 32(9), 1813–1822. doi:10.1162/jocn_a_01584

Tremblay, S., Rogasch, N. C., Premoli, I., Blumberger, D. M., Casarotto, S., Chen, R., … Daskalakis, Z. J. (2019). Clinical utility and prospective of TMS–EEG. Clinical Neurophysiology, 130(5), 802–844. doi:https://doi.org/10.1016/j.clinph.2019.01.001

Tröndle, M., Popov, T., & Langer, N. (2020). Decomposing the role of alpha oscillations during brain maturation. bioRxiv, 2020.2011.2006.370882. doi:10.1101/2020.11.06.370882

Uhlhaas, P. J., Roux, F., Rodriguez, E., Rotarska-Jagiela, A., & Singer, W. (2010). Neural synchrony and the development of cortical networks. Trends Cogn Sci, 14(2), 72–80. doi:10.1016/j.tics.2009.12.002

Voytek, B., & Knight, R. T. (2015). Dynamic network communication as a unifying neural basis for cognition, development, aging, and disease. Biol Psychiatry, 77(12), 1089–1097. doi:10.1016/j.biopsych.2015.04.016

Voytek, B., Kramer, M. A., Case, J., Lepage, K. Q., Tempesta, Z. R., Knight, R. T., & Gazzaley, A. (2015). Age-Related Changes in 1/f Neural Electrophysiological Noise. J Neurosci, 35(38), 13257–13265. doi:10.1523/JNEUROSCI.2332-14.2015

Wang, J., Barstein, J., Ethridge, L. E., Mosconi, M. W., Takarae, Y., & Sweeney, J. A. (2013). Resting state EEG abnormalities in autism spectrum disorders. Journal of Neurodevelopmental Disorders, 5(1), 24. doi:10.1186/1866-1955-5-24

Waschke, L., Donoghue, T., Fiedler, L., Smith, S., Garrett, D. D., Voytek, B., & Obleser, J. (2021). Modality-specific tracking of attention and sensory statistics in the human electrophysiological spectral exponent. bioRxiv, 2021.2001.2013.426522. doi:10.1101/2021.01.13.426522

Wilkinson, C. L., & Nelson, C. A. (2021). Increased aperiodic gamma power in young boys with Fragile X Syndrome is associated with better language ability. Molecular Autism, 12(1), 17. doi:10.1186/s13229-021-00425-x

Wolters, C. H., Anwander, A., Tricoche, X., Weinstein, D., Koch, M. A., & MacLeod, R. S. (2006). Influence of tissue conductivity anisotropy on EEG/MEG field and return current computation in a realistic head model: a simulation and visualization study using high-resolution finite element modeling. Neuroimage, 30(3), 813–826. doi:10.1016/j.neuroimage.2005.10.014

Yeo, I. K., & Johnson, R. A. (2000). A new family of power transformations to improve normality or symmetry. Biometrika, 87(4), 954–959. doi:10.1093/biomet/87.4.954

Yuval-Greenberg, S., Tomer, O., Keren, A. S., Nelken, I., & Deouell, L. Y. (2008). Transient induced gamma-band response in EEG as a manifestation of miniature saccades. Neuron, 58(3), 429–441. doi:10.1016/j.neuron.2008.03.027

