## Supplemental_Material for "Periodic and Aperiodic Neural Activity Displays Age-Dependent Changes Across Early-to-Middle Childhood"

### SUPPLEMENTARY MATERIALS AND METHODS

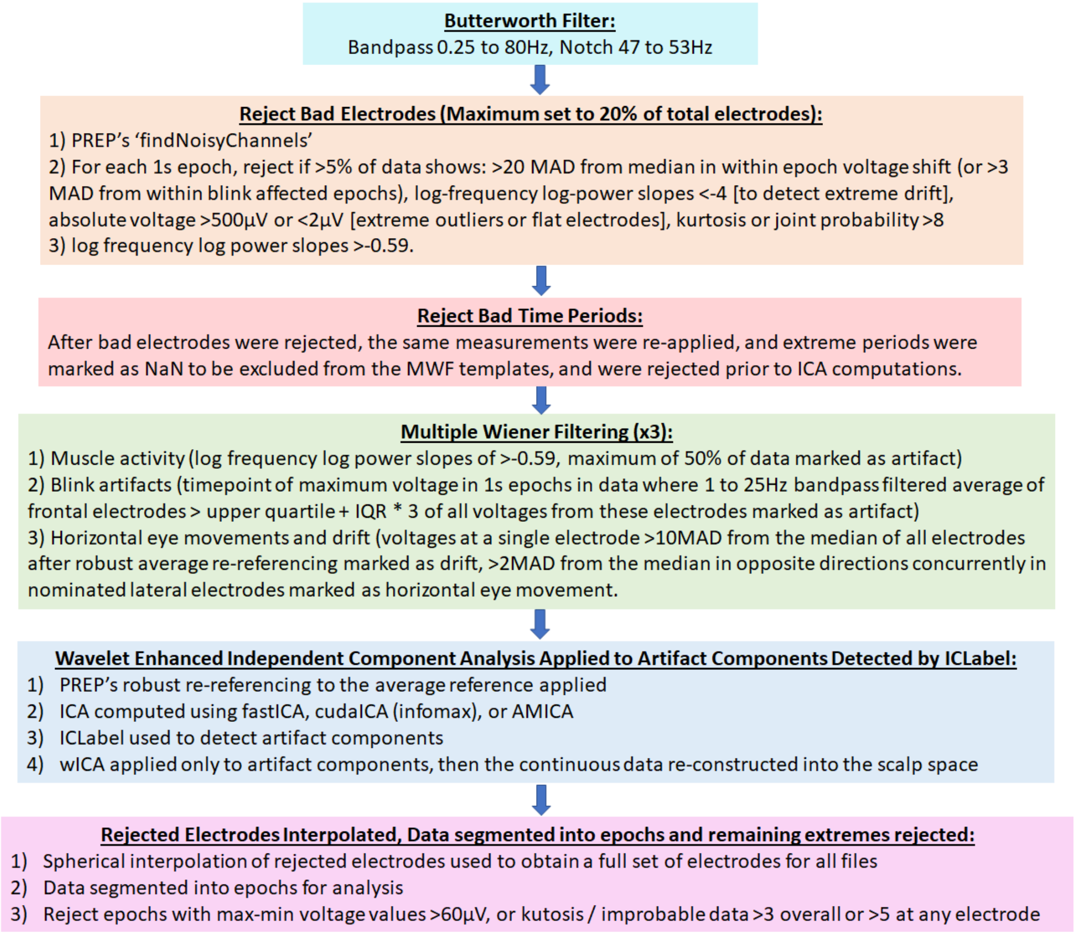


**Figure S1.** Overview of the typical steps involved in the Reduction of Electrophysiological Artefacts (RELAX) pipeline which was utilised to pre-process the resting-state EEG data (from Bailey et al., 2021).

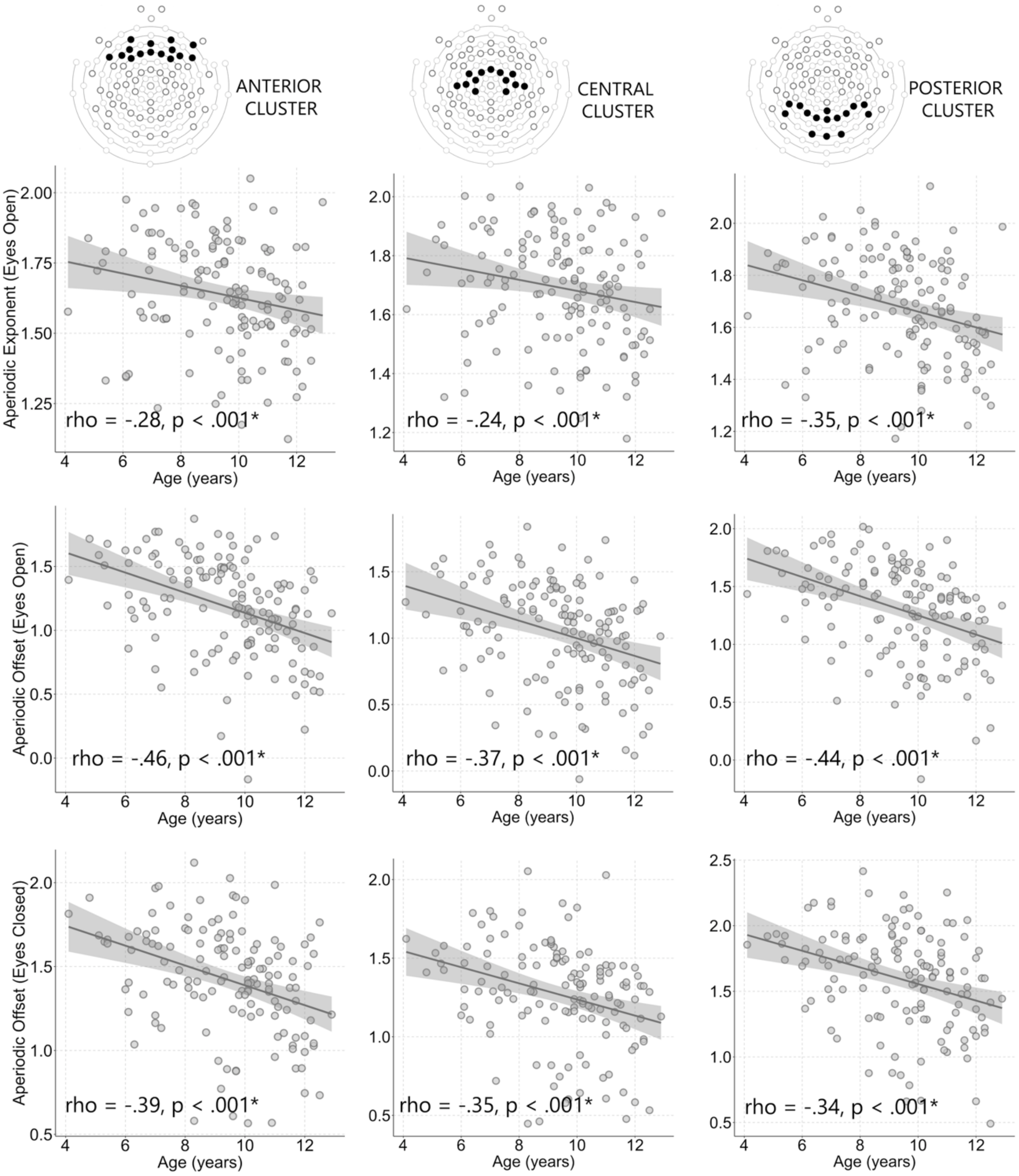


**Figure S2:** Correlations between age and either aperiodic exponent (eyes open), or offset (eyes open, eyes closed) for electrode clusters taken from anterior, central, and posterior regions. Regardless of the specific cluster used, there was a negative association between age and either aperiodic exponent, or offset.

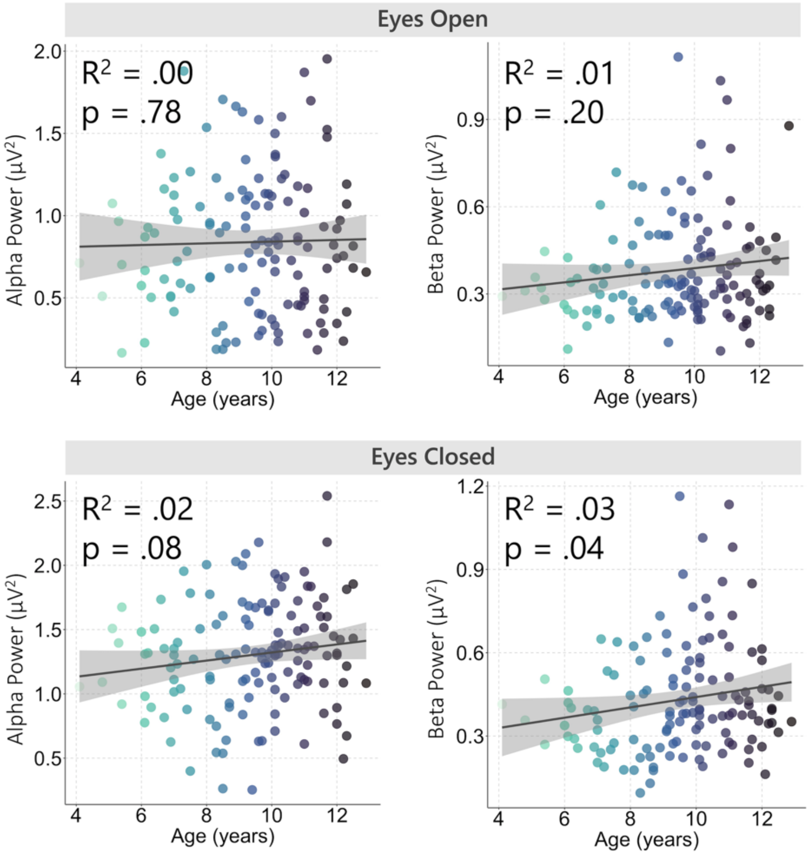


**Figure S3:** Scatter plots of spectral power in relation to age for the alpha and beta frequency bands. Significance values from the regression analyses are shown. Age was unable to significantly predict power within either the alpha or beta frequency ranges (after Bonferroni correction) for either the eyes open, or eyes closed data.

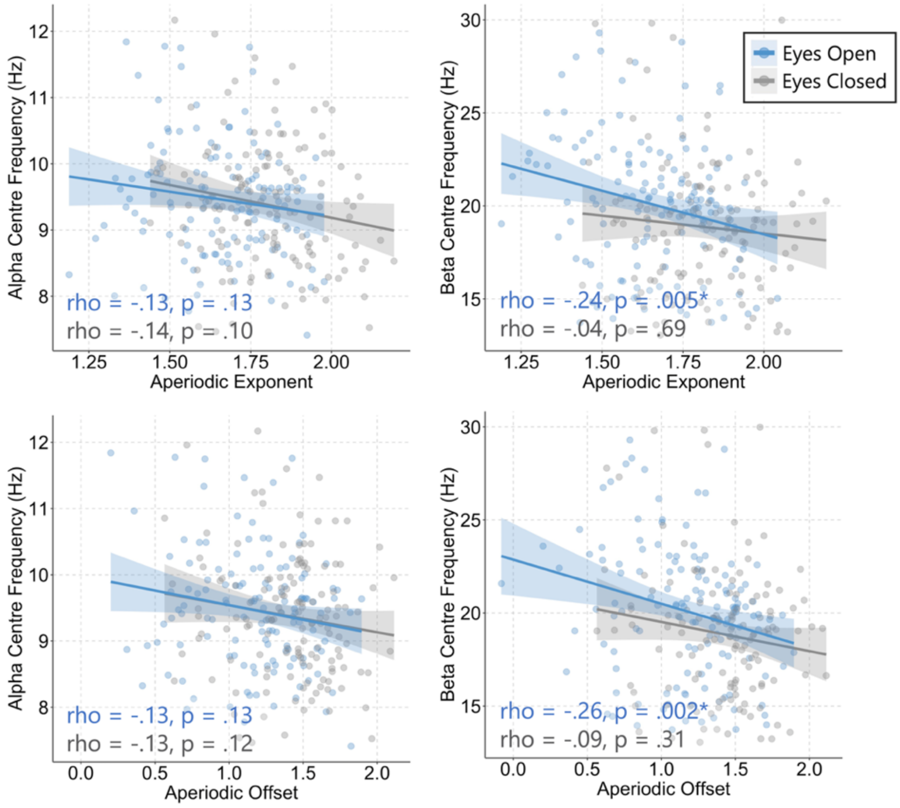


**Figure S4:** Correlations between alpha and beta centre frequency and aperiodic exponent (upper panel), as well as aperiodic offset (lower panel). Significant associations were found between beta centre frequency and aperiodic exponent and offset for the eyes open conditions only. Asterisk denotes significant result after Bonferroni correction.

**References**

Bailey, N. W., Biabani, M., Hill, A. T., Rogasch, N. C., McQueen, B., & Fitzgerald, P. B. (2021). Introducing RELAX (the Reduction of Electrophysiological Artifacts): A fully automatic pre-processing pipeline for EEG data. *In Preparation*.
